# Reinterpretation of *Bakiribu waridza* from the Romualdo Formation (Lower Cretaceous) of Brazil: a fish not a pterosaur

**DOI:** 10.64898/2025.12.10.693067

**Authors:** David M. Unwin, Roy E. Smith, Samuel Cooper, David M. Martill

## Abstract

Fragmentary remains of fossil vertebrates from the Romualdo Formation (Lower Cretaceous) of Brazil, preserved in association with two fish were interpreted as two individuals of a new genus and species of ctenochasmatine pterosaur, *Bakiribu waridza* Pêgas *et al*., 2025. Comparison with a range of fossils from the same geological unit show that the remains represent the gill arch apparatus of a large actinopterygian fish. This reinterpretation, which mirrors a comparable event in the mid twentieth century when the remains of a supposed ctenochasmatine pterosaur, *Belonochasma aenigmaticum*, were reidentified as part of the gill apparatus of a fish, does not affect our current understanding of the evolutionary history of pterosaurs.

## Introduction

Pêgas *et al*.^1^ recently described a new genus and species of ctenochasmatine pterosaur, *Bakiribu waridza*, from the Lower Cretaceous Romualdo (= Santana) Formation of Brazil, a well-known fossil Lagerstätte that has yielded a diverse assemblage of pterodactyloid pterosaurs^2^. The multiple fragments of this pterosaur, purportedly representing two individuals and preserved together with two fish in a single nodule (MCC 1271.1-V / MPSC 7312), were interpreted by Pêgas *et al*. as a ‘regurgitalite’.

Comparison with fossil vertebrates from the Santana Formation revealed features which indicate that the holotype (MCC 1271-Va/MPSC 7312a) and the paratype (MCC 1271-Ve/MPSC 7312e) of *Bakiribu waridza* are the remains of a large actinopterygian fish, and not a pterosaur.

## Methodology

### Data collection and materials

Data for this study was drawn from fossil material of *Ctenochasma elegans* (SNSB-BSPG 1935.I.24, SNSB-BSPG 1920.I.57, JME SoS 2179, JME SoS 2476, SMNS 8180) and *Belonochasma aenigmaticum* (SNSB-BSPG 1938 I 87). Approaches adopted include visual inspection, binocular microscopy and high-definition ultra-violet fluorescence photography. Additional data on *Bakiribu waridza* (MCC 1271-V/MPSC 7312) was collected from the original paper and supplementary digital materials published by Pêgas *et al*.^1^. Data on the gill arch morphology of amiid fish was assembled from the monographic study by Grande and Bemis^3^.

### Institutional abbreviations

**JME SoS**, Jura-Museum, Eichstätt, Bavaria, Germany; **MCC**, Museu Câmara Cascudo, Natal, Rio Grande do Norte, Brazil; **MPSC**, Museu de Paleontologia Plácido Cidade Nuvens, Santana do Cariri, Ceará, Brazil; **SMNS**, Staatliches Museum für Naturkunde, Stuttgart, Baden-Württemberg, Germany; **SNSB-BSPG**, Bayerische Staatssammlung für Paläontologie und Geologie, Munich, Bavaria, Germany.

### Reinterpretation of MCC 1271-Va/MPSC 7312 as a fish

The elongate bony elements interpreted by Pêgas *et al*.^1^ as fragments of the rostrum and mandibular symphysis of two pterosaurs are reinterpreted here as the gill arch apparatus of a large actinopterygian fish (Figures 1-3). Identifiable elements include a pair of robust anterior ceratohyals, closely comparable in form to those of *Cratoamia*^4^ each articulated with a hypohyal. Posteriorly the hypohyals are in contact with a basibranchial, a midline element to either side of which are the first two pairs of hypobranchials. This configuration (Figure 3A) matches almost exactly that described for *Cratoamia* (ref. 4, fig 6; Figure 2B). The remaining hypobranchials are displaced and, in most cases, incomplete (Figure 3A).

**Figure 1.**
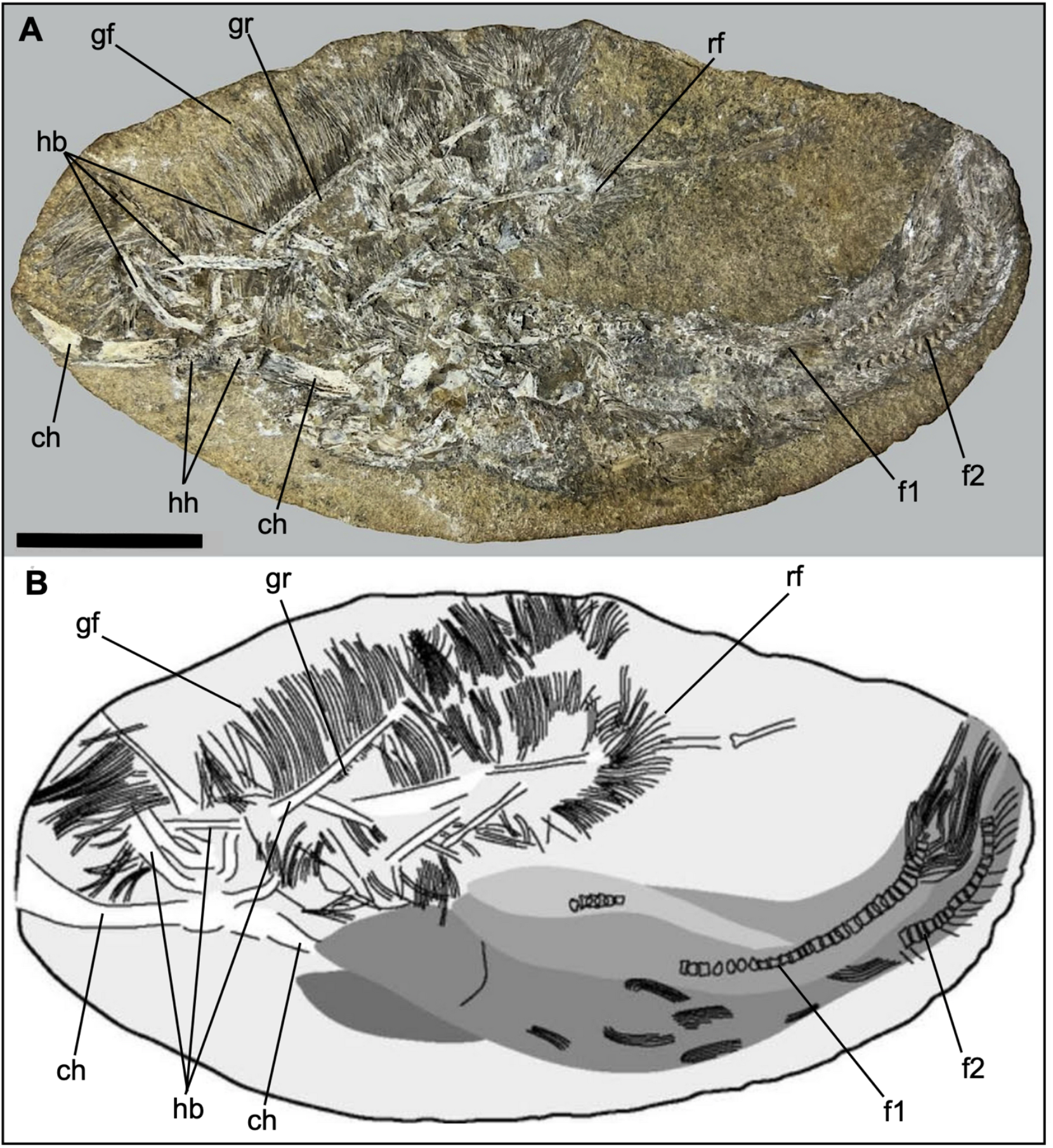
Reinterpretation of MPSC 7312 based on Pêgas *et al*., 2025. (A) Fossil remains. (B) Sketch of associated fish remains including gill arch apparatus of a large amiid and two examples of *Tharrhias araripis* (?). Abbreviations: **ch**, ceratohyal (cf. *Amia*); **f1**, fish 1; **f2** fish 2; **gf**, gill filaments; **gr**, gill rakers; **hb**, hypobranchial; **hh**, hypohyal; **rf**, reflexed hypobranchial. Scale bar = 50 mm.

**Figure 2.**
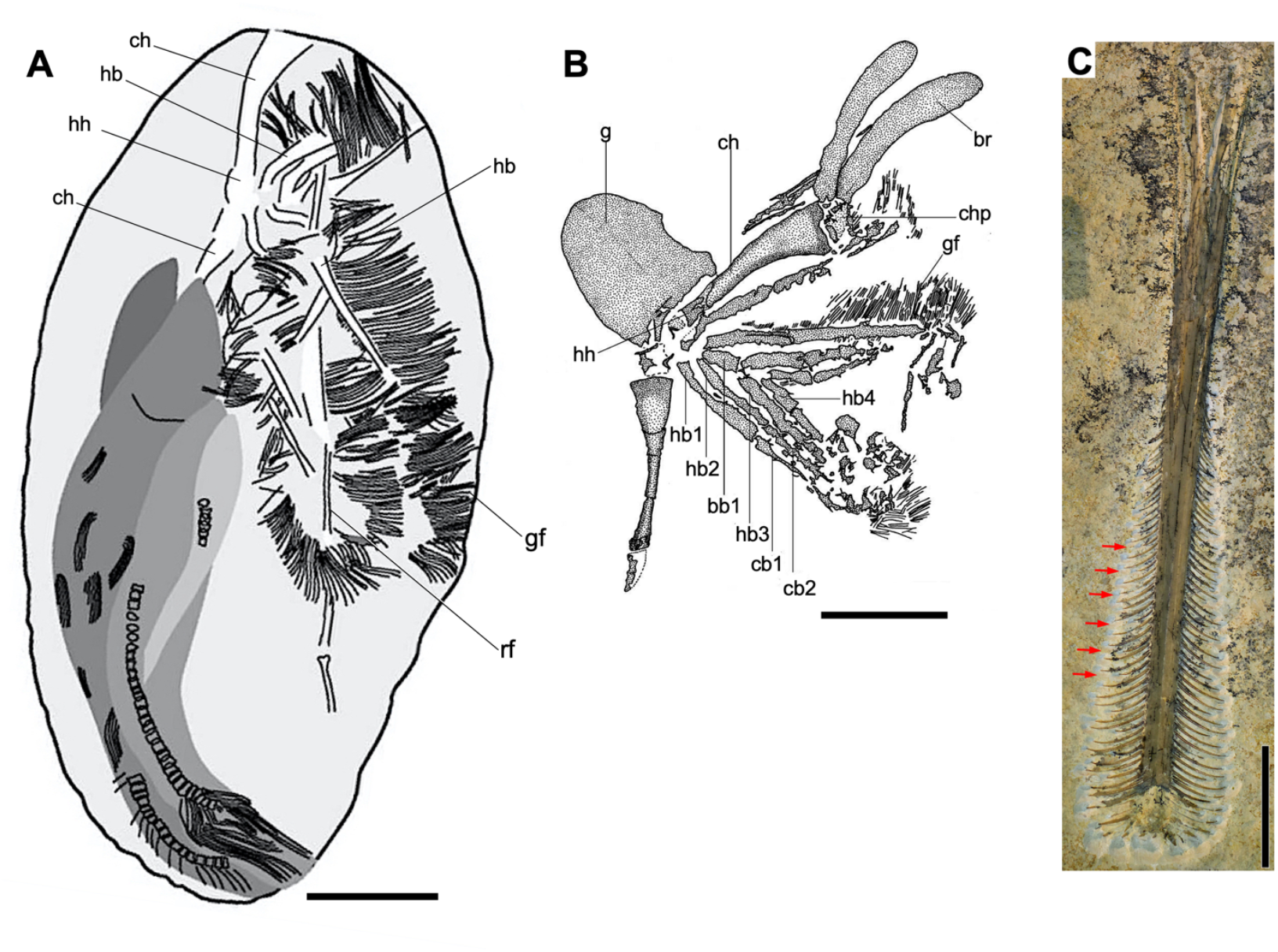
Comparison of MPSC 7312 with the gill arch of a fish and the rostrum of a ctenochasmatine pterosaur. (A). Line drawing of MPSC 7312 rotated vertically to facilitate comparison. (B). *Cratoamia gondwanica* (UERJ-PMB89), Crato Formation (Lower Cretaceous), Araripe, Brazil. Gular plate, hyoid arch and gill arches. (C). *Ctenochasma elegans* (JME SoS 2476), Altmühltal Formation (Upper Jurassic: Tithonian), Solnhofen, Germany. Rostrum with complete dentition in dorsal view. Arrows indicate immature teeth. Abbreviations: **bb**, basibranchial; **cb**, ceratobranchial; **ch**, ceratohyal; **chp**, posterior ceratohyal; **gf**, gill filaments; **hb**, hypobranchial; **hh**, hypohyal; **rf**, reflexed portion of hypobranchial. Scale bar = 25 mm. (A) Compiled from Pêgas *et al*., 2025; (B) reproduced from Brito *et al*., 2008, fig. 6); (C) Photo courtesy of R. Belben.

**Figure 3.**
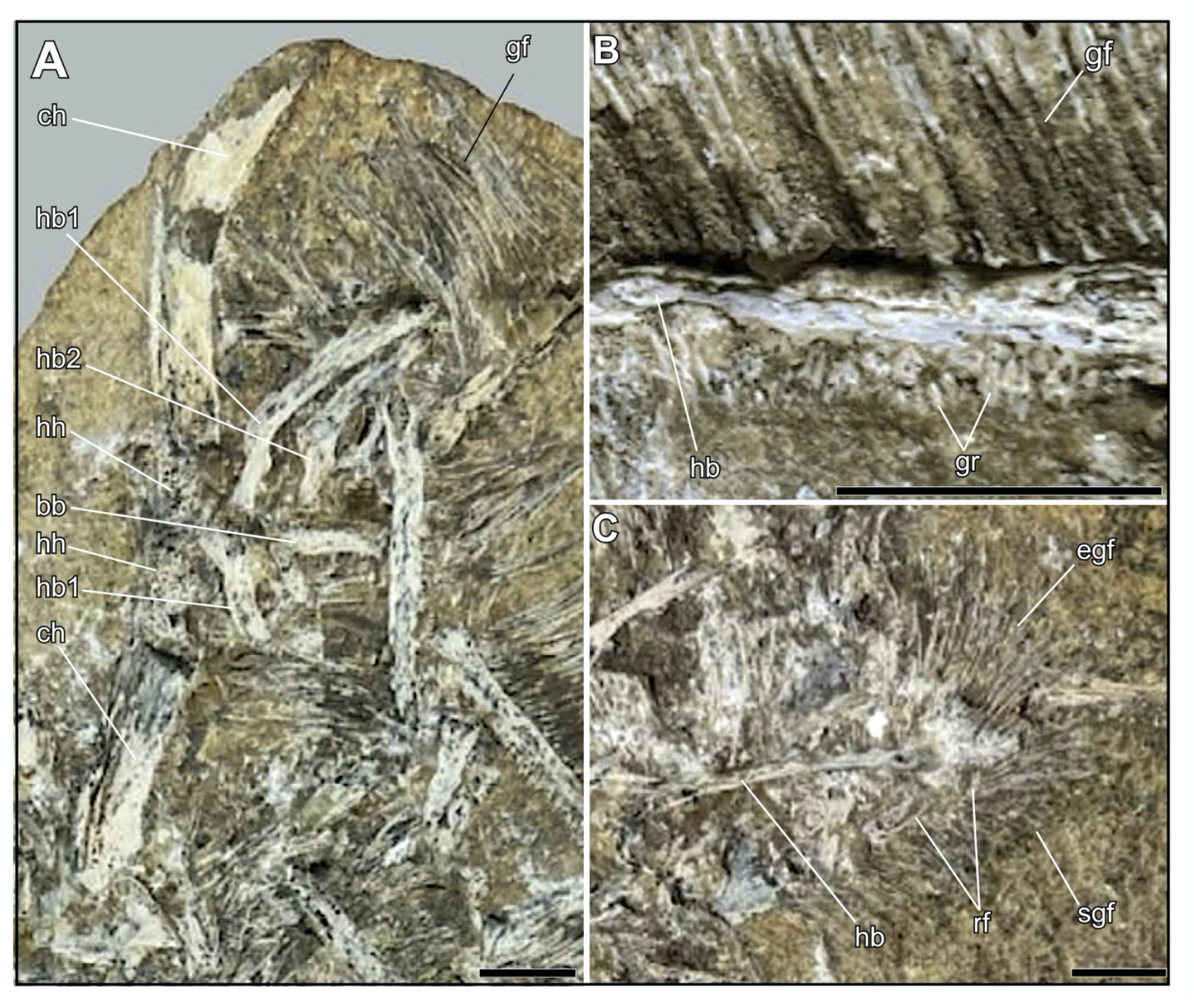
Anatomical details of the gill arches of MCC 1271-Va, e/MPSC 7312a, e. (A) Partially articulated remains of the gill arch assembly with a basibranchial, ceratohyals, hypohyals and hypobranchials #1 and #2 in natural association. (B) Gill rakers. (C) A reflexed hyobranchial element with elongate gill filaments proximally and relatively short gill filaments distally. Abbreviations: **bb**, basibranchial; **ch**, ceratohyal; **chp**, posterior ceratohyal; **egf**, elongate gill filaments; **gf**, gill filaments; **gr**, gill rakers; **hb**, hypobranchial; **hh**, hypohyal; **rf**, reflexed portion of hypobranchial; **sgf**, relatively short gill filaments. Scale bar = 25 mm. (Compiled from Pêgas *et al*., 2025).

As in *Cratoamia* the hypobranchials are elongate elements that taper distally. Along one margin each hypobranchial bears numerous tightly packed highly elongate gill filaments oriented perpendicular to the long axis of the hypobranchial. In well preserved examples, small, short gill rakers fringe the other margin of the hypobranchial (ref. 1, fig. 4A; Figure 3B).

The majority of the preserved portions of the hypobranchials and associated gill filaments appear to represent basal portions of the hypobranchial. One example (Figure 3C) appears to be more complete preserving the distal portion of the hypobranchial though only fragments of this remain. This part of the hypobranchial is sharply reflected toward the base of the structure, exhibiting a hook-shaped morphology, with the gill elements fanning out around the external margin of the hook. Proceeding toward the tip of the hypobranchial these filaments exhibit a marked decline in length. This hypobranchial gill filament assembly is strikingly similar to that observed in *Amia*^5^ and an actinopterygian from the Upper Jurassic Solnhofen Limestones of Southern Germany, described under the name *Belonochasma*^6,7^

### Comparison of MCC 1271-Va, e/MPSC 7312a, e with ctenochasmatine pterosaurs

The slender, elongate needle-like elements interpreted by Pêgas *et al*.^1^ as teeth, reinterpreted here as gill filaments, bear a superficial similarity to the dentition of ctenochasmatine pterosaurs^8^. They differ from the latter, however, in several important respects. First, in ctenochasmatines the dentition is symmetrically arranged on either side of the rostrum and the mandibular symphysis^9–11^ (Figure 2C). By contrast in MCC 1271-Va, e/MPSC 7312a, e the needle-like elements are almost entirely restricted to just one side of the supporting spar. Second, the teeth of ctenochasmatines are longest anteriorly and show a highly regular reduction in length posteriorly such that the posterior-most are less than 30% the length of the anterior-most elements^9^ (Figure 2C). By contrast, in MCC 1271-Va, e/MPSC 7312a, e filaments in the posterior-most positions are as long, or longer, than filaments in anterior positions^1^ (Figures 1, 2A) opposite to the condition in ctenochasmatines. Moreover, in *Bakiribu* some filaments are illustrated as exceptionally elongate (ref. 1, fig. 3A), a pattern never encountered in ctenochasmatid dentitions. Third, the dentition of ctenochasmatines invariably shows distinct patterns of tooth replacement with shorter replacement teeth regularly intercalated between longer, mature, fully erupted teeth^12^ (Figure 2C). There is no evidence of such patterns in the filaments of MCC 1271-Va, e/MPSC 7312a, e (Figures 1–2).

Fourth, the tooth crowns of *Bakiribu* are described as ‘subquadrangular’ in cross-section, a morphology consistent with gill filaments but not the teeth of ctenochasmatids which are circular or sub-circular in cross-section^13^.

In addition to the jaws and dentition Pêgas *et al*.^1^ briefly mention that: “An indeterminate metatarsal and an indeterminate pedal phalanx are also present, as are some indeterminate bone fragments.” We were unable to locate or identify these bones.

## Discussion

### Identification of MCC 1271-Va, e/MPSC 7312a, e

Not one of the skeletal elements preserved on the main slab or counterpart, exhibits a morphological feature that is uniquely pterosaurian. The filament-like structures, interpreted as teeth by Pêgas *et al*., show a superficial resemblance to the dentition of ctenochasmatines but lack key pterosaurian characteristics such as symmetry in the distribution of teeth along the jaw margins and distinctive patterns in size variation related to waves of tooth replacement, universally present in dentate pterosaurs and particularly well developed in ctenochasmatines^12^.

The morphology of the skeletal elements and filaments and the disposition of those bones that remain in articulation compare closely to elements of the gill arch of a large actinopterygian. The remains are too fragmentary to permit a more precise taxonomic assignment, but we note similarities between MCC 1271-Va, e/MPSC 7312a, e and amiids from the Lower Cretaceous of South America including *Calamopleurus*^14^ and *Cratoamia*^4^.

### Preservation and taphonomy

The reinterpretation of MCC 1271-Va, e/MPSC 7312a, e presented here is not consistent with the idea that this fossil represents a regurgitalite. The part and counterpart bear the remains of two small fish, likely *Tharrhias*. Pêgas *et al*.^1^ reported four examples of this fish but only two are evident in their illustrations. These fishes are fossilised in association with the incomplete remains of a much larger actinopterygian fish, likely an amiid, and represented by much of the gill arch complex.

There are two potential explanations for this association. (1) The small fish may have become lodged in the buccal cavity of the larger fish as several examples of the Santana Formation amiid *Calamopleurus* with smaller fish lodged in their gape are known^15^. (2) The association may simply be fortuitous, the concretion forming around the smaller fish and part of the larger fish while most of latter lay beyond the concretion boundary.

The alternative interpretation, that the fossil includes the fragmentary remains of the jaws of two individual pterosaurs and two complete fish, requires a much more complex taphonomic scenario and one without precedent in the fossil record. Furthermore, the explanation, that the fossil represents a regurgitalite does not account for the observation that the violence required to render the ‘jaws’ into multiple fragments is highly inconsistent with the preservation of the ‘teeth’ which show little or no displacement or damage.

### Palaeontological déjà vu?

This is not the first occasion on which the remains of the gill arches of a fish have been mistaken for a ctenochasmatid pterosaur. In 1939 Broili^6^ described a new vertebrate, *Belonochasma aenigmaticum*, in which gill arches and gill filaments were interpreted as the jaws and dentition of a ctenochasmatid. This specimen (BSPG 1938.I.87) was later reidentified as the gill arches of a fish^16^ with Mayr^7^ providing a detailed account based on a series of specimens in which more complete remains of the fish are preserved. Several examples figured by Mayr (ref. 7, figs 1–3) show clear similarity to MCC 1271-Va, e/MPSC 7312a, e.

### Implications of the reidentification of MCC 1271-Va, e/MPSC 7312a, e for pterosaur evolutionary history

Prior to the publication of *Bakiribu waridza*, concluded here to be a fish, ctenochasmatine pterosaurs were unreported from the Lower Cretaceous of Brazil. *Cearadactylus atrox* Leonardi and Borgomanero, 1985^17^, interpreted by Unwin^18^ as a ctenochasmatid^19^, but lost during the fire that destroyed the Museu Nacional Brazil in 2018^2^, was reinterpreted as an ornithocheirid following the demonstration that the fragmentary remains of the skull and jaws had been inaccurately reconstructed^20^. *Unwindia trigonus* Martill, 2011^21^, while seemingly not a ctenochasmatine, appears to be a lonchodectid and thus a member of Ctenochasmatoidea^2, 8^ (= Archaeopterodactyloidea of some workers).

The reinterpretation of *Bakiribu waridza* as a fish does not significantly change our understanding of pterosaur evolutionary history. Prior to this study, ctenochasmatines, represented, for example, by *Pterodaustro*^22^, were already known to be present in South America, persisting there until at least the end of the Early Cretaceous^23^. The continued absence of ctenochasmatines sensu stricto from either the Crato or Santana Formation contrasts with their complete dominance of the ‘Loma del *Pterodaustro*’ deposits in Argentina^23^ suggesting that ecology was a primary control on the patterns of distribution seen in the pterosaur fossil record.

## Conclusions

MCC 1271.1-V/MPSC 7312 is a typical fish bearing Romualdo concretion and not a regurgitalite. *Bakiribu waridza* is a fish, not a pterosaur. In that MCC 1271-Va, e/MPSC 7312a, e appears to consist of indeterminate remains of an actinopterygian fish, possibly an amiid, the name *Bakiribu waridza* should be treated as a *nomen dubium*.

## Acknowledgements

We are grateful to Oliver Rauhut and Markus Moser (Bavarian State Collection for Paleontology and Geology, Munich, Germany); Valentina Rosina and the late Martin Röper (Bürgermeister-Müller-Museum, Solnhofen, Germany); Christina Ifrim and Andreas Hecker (Jura-Museum, Eichstätt, Germany); Georg Bergér (Museum Bergér, Harthof, Eichstätt, Germany); Erin Maxwell and Rainer Schoch (State Museum of Natural History, Stuttgart, Germany) and Mike Day (Natural History Museum, South Kensington, UK) for facilitating access to specimens in their collections. We thank Rab Smyth for commenting on earlier versions of the manuscript.

## Author contributions

Conceptualisation: D.M.U and D.M.M.; methodology: D.M.U., D.M.M., R.E.S and S.C.: investigation and analysis: D.M.U., D.M.M., R.E.S and S.C; illustrations: D.M.U.; writing: D.M.U., D.M.M., R.E.S and S.C.

## Competing interests

The authors declare no competing interests.

## Additional information

Correspondence and requests for materials should be addressed to D.M.U.

## Notes

### Competing Interest Statement

The authors have declared no competing interest.

